# Systematic Comparison of 10X Genomics and Parse Single Cell RNA Technologies across PBMC and CD8^+^ TEMRA Cells

**DOI:** 10.64898/2026.09.02.748788

**Authors:** Alejandro Espinoza, Bruno J. de Andrade Silva, Matteo Pellegrini

## Abstract

Single-cell RNA sequencing technologies provide insights into gene expression at the cellular level, enabling detailed analysis of cellular heterogeneity. In this study, we systematically compared two scRNA-seq platforms,10X Genomics and Parse Biosciences, using human peripheral blood mononuclear cells (PBMCs) and terminally differentiated effector memory CD8^+^ T cells (TEMRAs). We identified significant differences in gene expression variability and platform-specific biases, such as ribosomal and mitochondrial gene capture. 10X has a bias for shorter genes, while Parse exhibited enhanced detection of longer transcripts. In CD8^+^ TEMRAs, the expression of key genes related to antimicrobial immune responses were underrepresented in Parse cells compared to 10X cells (e.g. *GNLY*, *PRF1* and *GZMB*). These findings underscore the need for careful selection of scRNA-seq platforms based on specific research objectives, as platform-specific biases can influence cell type identification and as well as mechanistic insights derived from gene expression data. Our results provide insights for selection of scRNA-seq experimental platforms in immunological studies.

## INTRODUCTION

Single-cell RNA sequencing^1^ (scRNA-seq) has revolutionized our understanding of cellular biology, enabling researchers to explore gene expression profiles at unprecedented resolution^2^. This technology has rapidly evolved, with new methods becoming available over the past few years, making it increasingly accessible to researchers across various fields^3,4^. scRNA-seq has been instrumental for the study of cellular heterogeneity in complex tissues, identifying rare cell populations, and elucidating developmental trajectories in various biological systems and diseases^5,6^.

The power of scRNA-seq lies in its ability to quantify gene expression within individual cells, overcoming the limitations of bulk RNA sequencing (bulk RNA-seq) methods that average signals across cell populations^7^. This granular view has proven particularly valuable in fields such as immunology^8^, neuroscience, and cancer research, where cellular diversity plays a crucial role in tissue function and disease progression^9,10^. However, scRNA-seq is not without limitations; it can be affected by technical challenges such as dropout events, where genes are not detected due to inefficient mRNA capture or amplification, potentially leading to biased or incomplete transcriptional profiles^8^. As new scRNA-seq technologies emerge, evaluating their performance against established methods is crucial for researchers to make informed decisions about which platform best suits their experimental needs. Droplet-based methods, which capture cells in oil droplets, are commonly used due to their high throughput and relative ease of use^11^. However, newer methods offer alternative approaches that may provide advantages in certain experimental contexts. Most recently, split-pool ligation sequencing, a well-based approach, has become commercially available, promising high-throughput capabilities with potential cost benefits^12^.

A recent study compared 10X Fixed RNA Profiling (FRP; also known as Flex Gene Expression) to alternative platforms, including HIVE CLX scRNA seq v1 (Honeycomb Technologies) and Evercode WT kit v2 (Parse Biosciences), finding that FRP provided superior performance in gene detection, cell clustering, and discrete cell type identification in PBMCs^13^. Another study, comparing 10X Genomics Chromium single-cell RNA sequencing, hereafter termed, 10X, and Parse Biosciences V2 split-pool ligation-based transcriptome sequencing, hereafter termed, Parse, highlighted that these platforms can yield different results in terms of gene detection sensitivity, cell type identification, and overall data quality^14^. This study utilized human peripheral blood mononuclear cells (PBMCs) to compare the platforms. Both technologies showed similar detection of genes across expression levels although the expression of shorter genes was higher in 10X. They identified similar cell types, but the Parse platform showed a decrease in T cell gene expression counts a T cell sub populations which remains unexplored.

CD8^+^ TEMRA cells (T effector memory cells re-expressing CD45RA) are a subset of memory T cells characterized by their CCR7-negative and CD45RA-positive phenotype. These cells are terminally differentiated CD8^+^ cytolytic T cells (CTLs), such that they have a distinct gene expression signature, including high expression of cytotoxicity-associated genes like granzymes, perforin and granulysin^15–17^. We have previously shown that CD8^+^ TEMRA cells are particularly enriched for tricytotoxic T lymphocytes (T-CTL), a T cell subset which simultaneously expresses granzyme B, perforin, and granulysin, making them a crucial component of the adaptive immune response against *Mycobacterium tuberculosis* (Mtb) and *Mycobacterium leprae* (mLEP)^15–16,18–20^. The intracellular staining for T-CTL markers precludes functional studies. However, the expression of surface receptors on CD8^+^ TEMRA cells, which are composed primarily of T-CTL, allows those cells to be isolated and functionally studied at the transcriptional level, which was previously precluded by the cell fixation required to label granzyme B, perforin, and granulysin. Thus, the investigation of CD8^+^ TEMRA cells presents a valuable opportunity to assess how different methodologies capture subtle variations within a relatively homogeneous population^15^. More specifically, in CD8^+^ TEMRA cells, the co-expression levels of *GZMB*, *GNLY* and *PRF1* genes directly affect different functional cell states.

Here we compared 10X Genomics Chromium 3’ single-cell RNA sequencing and Parse Biosciences V2 split-pool ligation-based transcriptome sequencing, using both a heterogenous cell population of PBMCs and a homogenous cell type, CD8^+^ TEMRAs, to understand differences in RNA expression capture in cell populations between platforms. By systematically evaluating these cells, we aimed to establish a comprehensive comparative framework for assessing the technical performance and transcriptional insights offered by these single-cell technologies. Our findings have important implications for researchers studying immune cell populations and highlight the need for careful consideration when selecting scRNA-seq platforms for specific research questions.

## RESULTS

### Quality control analysis of PBMCs

We generated six PBMCs libraries for 10X and Parse from three donors (D000, D252 and D394) with matched samples for each platform. PBMCs were collected for their accessibility, viability in single-cell assays, and their heterogeneous cell type populations making them useful for benchmarking. The single cell libraries were aligned (see Methods) and initial cell counts were 49,298 for 10X and 49,692 cells for Parse across the three donors. We processed the data using Seurat, and after normalization, we observed the average unique molecular identifiers (UMIs) per cell to be greater in 10X compared to Parse (Supplementary Fig 1A and B). The average number of transcripts captured per cell was significantly greater (Wilcoxon rank sum test with continuity correction, p-value < 2.2e-16) in Parse, compared to 10X (Fig 1A and B). 10X captured more mitochondrial gene transcripts per cell compared to Parse (Supplementary Fig C). We removed cells containing more than 20% mitochondrial genes per cell in both platforms (Supplementary Fig 1D). After down sampling and filtering to correct for library sequencing depth, we collected a similar number of unique genes (Supplementary Fig 1E), UMIs and mitochondrial gene percentage between 10X and Parse libraries. The final analysis used 37,809 10X cells and 37,529 Parse cells across three donors.

**Fig 1.**
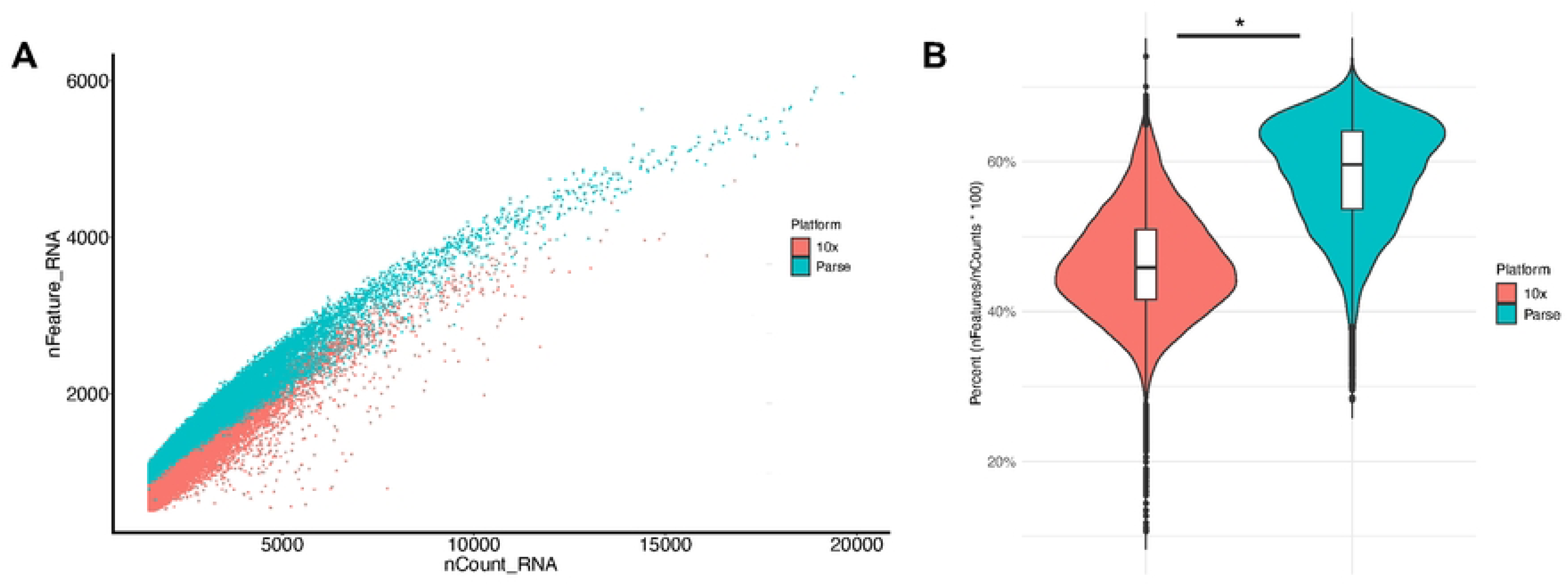
Comparison of PBMCs’ RNA expression profiles between 10X and Parse technologies. (A) Scatter plot showing the relationship between unique molecular identifiers (nCount_RNA) and number of unique genes per cell (nFeature_RNA). (B) Violin plots depicting the distribution of the percent of unique genes per UMI for each cell (Wilcoxon rank sum test with continuity correction, p-value < 2.2e-16).

### Transcriptomic differences in gene abundance captured due to gene length

We performed a systematic analysis of protein-coding genes by averaging the total counts of UMIs per gene for 10X and Parse and identified high concordance in gene expression (Pearson’s r = 0.904) in protein coding genes (Fig 2A). Although we observed a large correlation in gene expression between platforms, ribosomal coding genes were more abundantly expressed in 10X compared to Parse. 10X consistently yielded higher ribosomal gene average counts, suggesting a potential bias in capturing this abundant mRNA population. We calculated the percentage of ribosomal transcripts (both large and small subunits) per cell. Our analysis revealed a significant difference (p-val < 0.05) in ribosomal gene expression per cell between 10X (average, 5.58%) and Parse (average, 0.38%) platforms (Fig 2B), highlighting a substantial technical variation in ribosomal gene detection.

**Fig 2.**
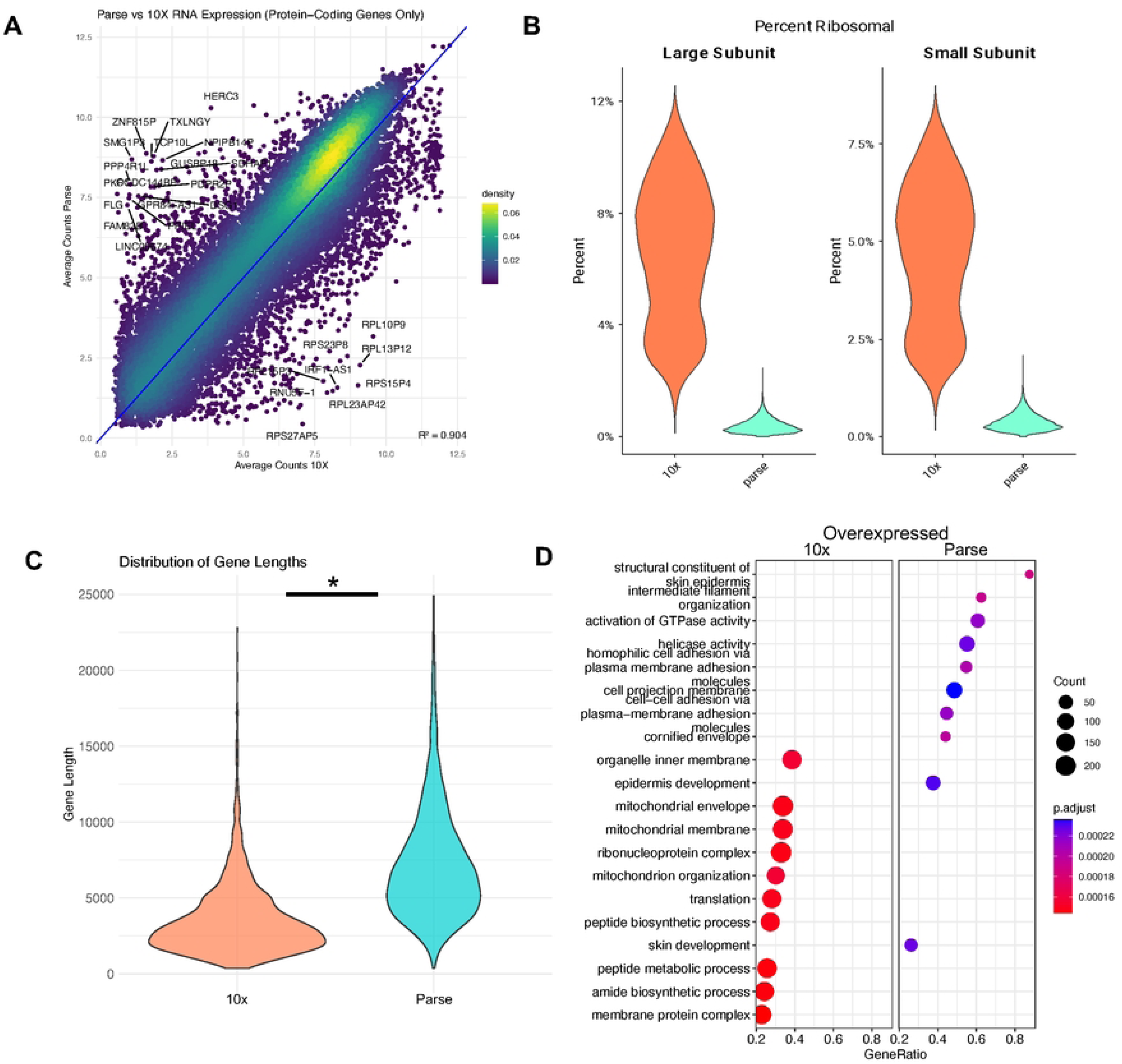
Comparison of 10X and Parse RNA expression data. (A) Scatterplot of average protein coding gene expression between 10X and Parse. (B) Violin plots showing percent ribosomal content across platforms. (C) Violin plots of distribution of overexpressed gene lengths between platforms. (D) Gene ontology enrichment analysis of overexpressed genes between 10X(left) and Parse (right).

To elucidate the underlying factors contributing to method-specific biases, we conducted differential gene expression analysis between platforms. Genes preferentially detected by Parse were longer, with transcript lengths exceeding 10X-detected genes by >1000bp, on average (Fig 2C). This observation suggests that Parse may offer enhanced sensitivity for longer transcripts, potentially due to differences in molecular capture efficiency. We compared the gene capture efficiency using bulk RNA-seq data from the PBMCs of seven healthy human individuals^19^. The average protein coding gene size in the bulk RNA-seq (4686.52) is closer to 10X (4674.52) than Parse (4769.76), although both methods share a similar distribution of gene lengths (Supplementary Fig 2A). Additionally, by comparing the weighted (log_10_(average gene expression * gene length)) scores of protein coding genes, we observed a similar transcript profile between Parse and 10X but not compared to bulk (Supplementary Fig 2B); we observed shorter genes to be more highly expressed in the bulk RNA-seq dataset.

Gene Ontology (GO) enrichment analysis of differentially expressed genes showed distinct functional profiles associated with each method (Fig 2D). 10X upregulated genes were enriched for fundamental cellular processes, including organelle organization, mitochondrial function, and translation. By contrast, upregulated genes in Parse showed significant enrichment for biological processes related to epidermal development, cell adhesion, and extracellular matrix organization.

### Unsupervised clustering identifies differences in immune cells

We performed unsupervised clustering on PBMCs from three shared donors (D000, D252, D394) using independent workflows for 10X and Parse datasets. To account for donor-specific batch effects, we applied Harmony integration separately to each platform. The resulting UMAP embeddings demonstrated consistent alignment of shared cell states across donors within each platform, indicating effective batch correction and comparable integration performance (Fig 3A). Stacked bar plots revealed that most clusters incorporated cells from all three donors, further supporting integration quality (Fig 3A).

**Fig 3.**
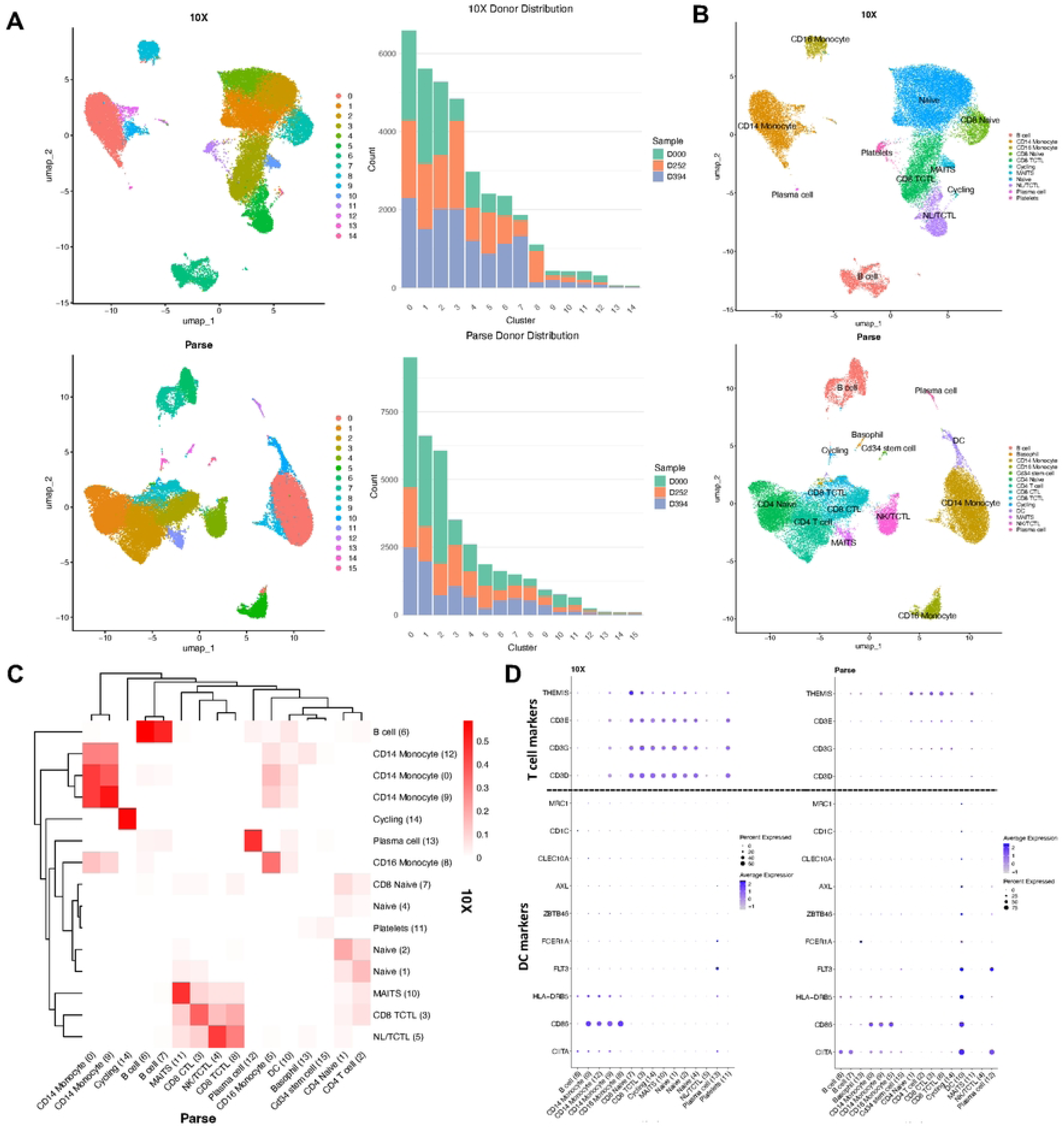
PBMC Cell Composition Captured by 10X and Parse. (A) UMAP of the integrated dataset colored by unsupervised clusters for 10X (top) and Parse (bottom). Adjacent stacked bar charts quantify donor contributions to each cluster. (B) Manually annotated major cell types visualized on the same UMAP embeddings for 10X (top) and Parse (bottom). (C) Heatmap of cell-type Jaccard overlap in 10X clusters (rows) compared to Parse (columns). (D) Dot plots depicting expression of canonical T-cell-sub-lineage and DC markers (rows) within inferred cell types (columns) in both platforms. Dot diameter encodes the proportion of cells expressing the gene, and color intensity reflects average scaled expression.

To assign biological identities to the clusters, we first calculated differentially expressed genes for 15 clusters in 10X and 16 clusters in Parse, then matched the top 100 genes per cluster to a curated PBMCs reference marker list derived from bulk RNA-seq data (see Methods). This approach reduced potential circularity in using single-cell-derived markers for annotation. Based on this strategy, we annotated 11 canonical immune cell types in 10X data and 13 in Parse data (Fig 3B), including T cells, B cells, NK cells, monocytes, and dendritic cells (DCs).

Both platforms captured expected PBMCs compartments, although differences were evident in representation of specific subpopulations. For instance, 10X showed increased clustering of CD8⁺ T cells and naive T cells, whereas Parse revealed rare populations such as basophils, DCs, and CD34⁺ hematopoietic progenitors (Fig 3B), which were not detected in 10X.

To quantify marker concordance, we computed the Jaccard similarity of top marker genes between 10X and Parse clusters (Fig 3C and Supplementary Fig 3). While most major cell types showed high agreement, T cells and DCs exhibited notably lower similarity scores, suggesting possible differences in detection sensitivity or resolution between platforms.

We further explored this discrepancy by comparing expression levels of canonical T cell markers (*CD3D*, *CD3E*, *CD3G*) and DC markers across platforms (Fig 3D). Parse cells showed better detection and expression of DC markers compared to 10X. In contrast, Parse captured reduced expression of *CD3* genes in comparison to 10X, implying lower capture efficiency or transcript recovery in T cell populations. To validate these findings using an orthogonal annotation strategy, we applied Azimuth, leveraging its human PBMCs reference dataset to perform automated cell type annotation (Supplementary Fig 4A and B). This enabled a hierarchical classification of immune cell identities and allowed cross-platform comparisons at multiple levels of granularity. At the Level 1 (L1) classification, we observed broadly consistent distributions of major immune populations, including B cells, NK cells, monocytes, and dendritic cells (Supplementary Fig 4C). However, the relative proportions of T cells and monocytes differed between platforms, with 10X recovering more T cells and Parse capturing a higher fraction of monocytes, possibly due to platform-specific biases in cell capture or lysis sensitivity.

At the finer-grained Level 2 (L2) resolution, more nuanced differences emerged, particularly among T cell subsets (Supplementary Fig 4D). The 10X platform captured higher frequencies of CD4⁺ effector memory (TEM), CD8⁺ central memory (TCM), and regulatory T cells (Tregs), consistent with its enhanced transcript recovery in these populations. In contrast, Parse displayed a pronounced enrichment of double-negative T cells (dnT), which may reflect differential sensitivity to surface marker dropout or altered cluster boundaries during integration. CD4⁺ TCM, CD8⁺ TEM, and NK cell frequencies were relatively well conserved between platforms, suggesting some degree of consistency in profiling these lineages.

### CD8^+^ TEMRA cells’ immune gene expression differs between platforms

To compare the two approaches on a less complex mixture of cell types, we conducted a comparative analysis of single-cell transcriptomics data generated from both 10X and Parse platforms to elucidate platform-specific differences in cell type identification and gene expression patterns from CD8^+^ TEMRA cells sorted from PBMCs (donor 302). We again performed down sampling of UMIs to ensure comparable UMIs between platforms for robust statistical comparisons. We validated the CD8^+^ TEMRA sorted cells (Supplementary Fig 5) by mapping the expression of PTPRC and noting the lack of expression of CCR7 genes in both platforms (Supplementary Fig 6). We collected a total of 2800 and 2915 cells for 10X and Parse platforms, respectively (Supplementary Table 1).

To evaluate cross-platform concordance in gene expression profiles, we first compared average log-normalized protein-coding gene expression across matched samples profiled by 10X and Parse. We observed a strong linear correlation between platforms (R² = 0.804), confirming broad agreement in transcript quantification despite technical differences (Fig 4A). Observing a subset of genes exhibited platform-specific biases, including overrepresentation of certain ribosomal and mitochondrial transcripts in Parse data.

**Fig 4.**
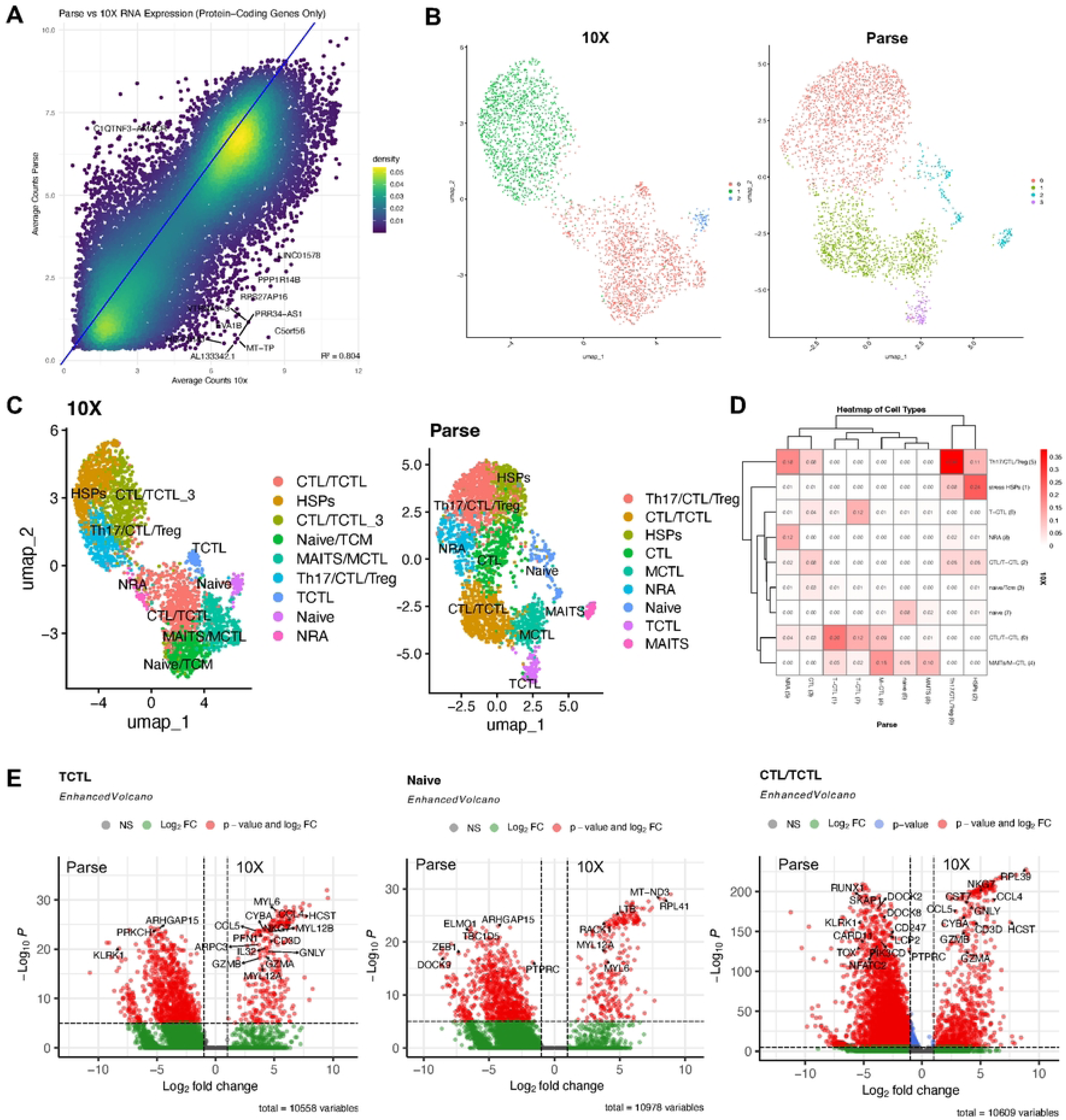
Cross-platform benchmarking of CD8⁺ TEMRA cells profiled by 10X and Parse. (A) Density scatterplot of mean log normalized counts per platform. (B) UMAP plot clustering of 10X (left) and Parse cells (right). (C) The same embeddings colored by curated cell-type assignments show preservation of major TEMRA sub-states. (D) Heat map of cell-type Jaccard similarity between 10X clusters (rows) compared to Parse (columns). (E) Volcano plot of differentially expressed protein coding genes between 10X (right) and Parse (left) in TCTL, Naive, and CTL/TCTL cell types, respectively.

Next, we performed unsupervised clustering using Seurat’s Louvain algorithm at a resolution of 0.5, revealing conserved cellular architecture across platforms (Fig 4B). Three major clusters were detected in 10X, whereas Parse yielded a more granular cluster structure, including subpopulations absent or merged in 10X. To resolve these populations, we manually annotated clusters based on a curated reference list of marker genes for CD8⁺ T cell subsets^15,20,21^. Annotated UMAP embeddings demonstrated preservation of major effector memory subtypes, including tri-cytotoxic T lymphocytes (TCTLs; *GZMB*, *PRF1* and *GNLY* co-expression), cytotoxic T lymphocytes (CTL)/TCTL hybrids, naive-like, MAIT, and Th17/Treg-like populations (Fig 4C and Supplementary Fig 7A).

To assess inter-platform cluster concordance, we calculated Jaccard similarity scores between the top 100 differentially expressed genes for each matched cluster. Overall, we observed poor overlap across platforms, with similarity scores below 50% in all cases (Fig 4D and Supplementary Fig 7B), indicating limited agreement in the most enriched transcripts defining each cluster.

To pinpoint genes contributing to these discrepancies, we performed pairwise differential expression analysis across three conserved subpopulations (TCTLs, Naive, and CTL/TCTL) between platforms (Fig 4E). We identified both conserved and platform-enriched genes. Shared markers such as *GZMH*, *PRF1*, and *CCL5* were robustly detected in both platforms, while platform-specific expression of genes such as *GNLY, GZMB, RUNX3*, *ELMO1*, *TRDC*, and *MYL12B* may reflect technical capture biases or differences in transcript isoform detection sensitivity in 10X cells.

To better understand the biological implications of platform-specific transcriptional differences, we investigated the expression of T cell-associated genes and pathways. We performed differential expression analysis between 10X and Parse cells. Comparing GC content between significantly upregulated markers indicated a slight bias in 10X GC content capture (p < 7.37e-9, Fig 5A). Further analysis revealed that genes preferentially detected by Parse tended to have longer total exon lengths (p < 2.22e–16, Fig 5B), implicating transcript length as a potential contributor to platform-specific bias in transcript capture.

**Fig 5.**
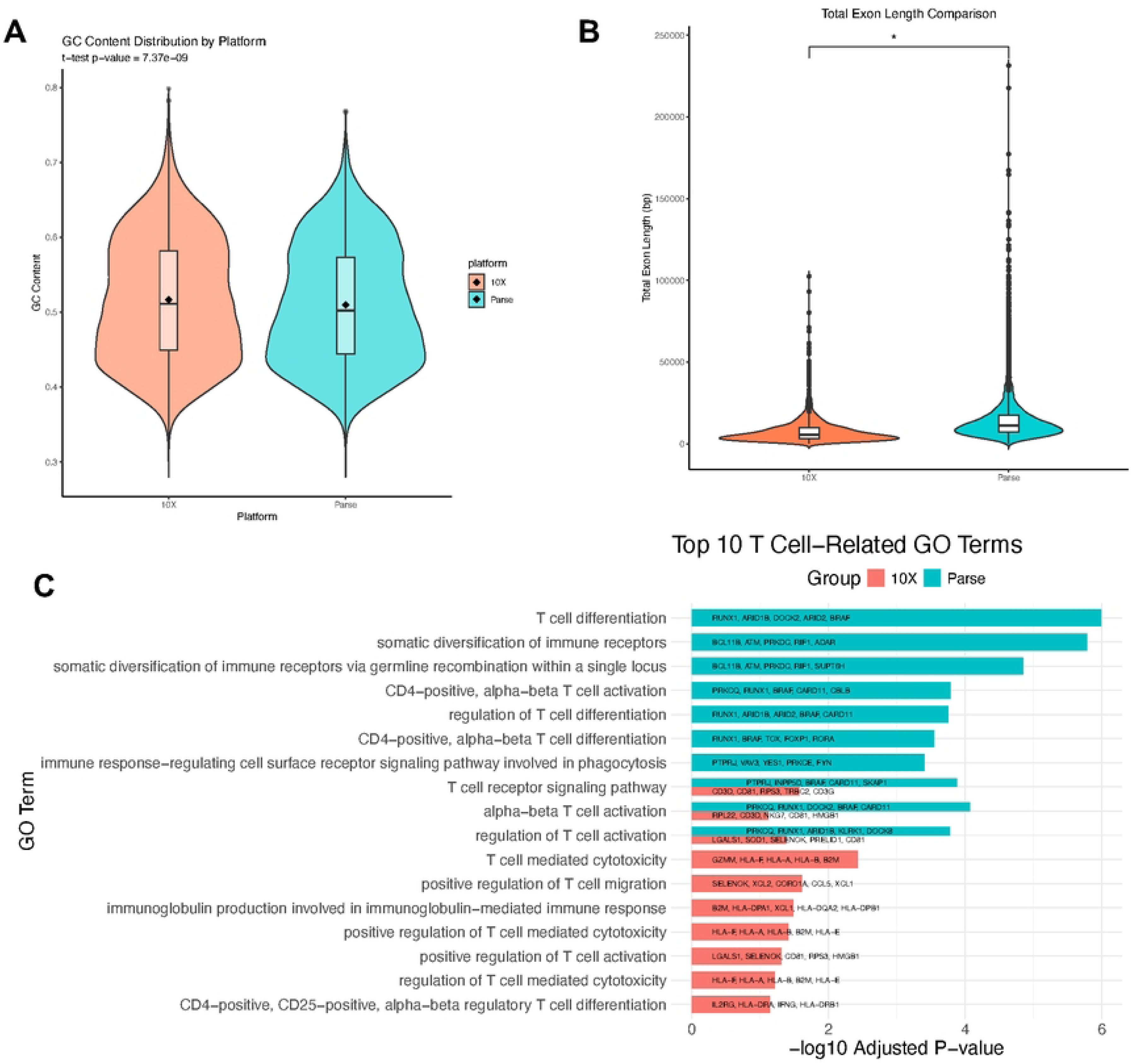
Platform-specific transcriptional signatures and T cell-associated pathways. (A) Gene expression profiles of known TEMRA genes. (B) Violin plots of distribution of overexpressed gene lengths between platforms (p value < 2.22e-16). (C) Enriched GO terms for T cell processes (top 10, -log10 adjusted P-value).

Gene ontology (GO) enrichment analysis of platform-enriched genes highlighted strong T cell-related signatures, such as “T cell differentiation,” “T cell activation,” and “T cell receptor signaling”, in 10X but not Parse (Fig 5C). These findings suggest a broader functional deficit in T cell profiling with Parse, possibly driven by chemistry or protocol-level constraints.

To validate these findings using an orthogonal annotation strategy, we applied Azimuth, leveraging the PBMC dataset. UMAP embeddings demonstrated all clusters to be mainly comprised of CD8 TEM cells in 10X and NK cells in Parse (Supplementary Fig 8A). To explore this at the single-gene level, we compared the expression of canonical T cell and cytotoxicity markers across platforms. Parse consistently showed reduced expression of *CD3D*, *CD3E*, *IL7R*, *CD8A*, *NKG7*, *GNLY*, *PRF1*, and *GZMB*, key identifiers of effector CD8⁺ T cells and TEMRA subsets (Supplementary Table 2). Cell-type annotations further revealed that while 10X classified the majority of CD8⁺ cytotoxic cells as CD8 TEM, Parse disproportionately labeled these cells as NK cells (Supplementary Fig 8B). This reclassification likely results from a dropout of CD3 and CD8 transcripts in Parse data, skewing the marker profile toward genes shared with cytotoxic NK cells such as *GZMB* and *GNLY*. Previously, it was demonstrated that CD8⁺ TEMRA cells are primarily composed of cytotoxic T cells, including TCTL subtypes^15,16,20^. To further characterize TEMRA heterogeneity, we scored cells using curated gene modules derived from bulk RNA-seq datasets^15^ for mono-, di-, and tri-cytotoxic T cells (MCTL, DCTL, TCTL) as well as MAIT cells, using Seurat’s ModuleScore() function^22^. 10X-derived cells exhibited significantly higher module scores compared to Parse, indicating greater enrichment of bulk-defined cytotoxic gene signatures (Supplementary Fig 9A and B). Specifically, 10X showed positive mean scores in three of the four subtypes (MCTL1: 0.022; TCTL1: 0.0437; MAIT1: 0.00007), while Parse exhibited negative scores across all subtypes, including MCTL1 (−0.0581) and TCTL1 (−0.0340), suggesting weaker cytotoxic signature capture.

Subtype classification based on these module scores revealed that 53% of 10X-derived cells were annotated as TCTLs, compared to only 19% in Parse (Supplementary Fig 9C). This reduced representation in Parse is lower and may reflect platform-specific limitations in detecting functionally distinct cytotoxic T cell states. These results support the conclusion that the 10X platform more accurately captures CD8⁺ TEMRA functional diversity.

## DISCUSSION

In this study, we performed a comprehensive comparison of single-cell RNA sequencing (scRNA-seq) platforms,10X Genomics and Parse Biosciences, using matched human PBMCs samples. While global transcriptomic profiles were highly concordant (R² > 0.9), we uncovered platform-specific differences in the resolution of immune cell subtypes, particularly CD8⁺ TEMRA cells. Across multiple analyses, 10X demonstrated superior sensitivity in capturing short, lowly expressed transcripts, including key immune effectors such as *GNLY*, *PRF1*, and *GZMB*, which were detected at significantly higher levels (average log₂ fold change > 2) compared to Parse. These differences have direct implications for studying human immune responses, as these genes are central to cytotoxic function and host defense.

When analyzing cell subtypes, both platforms recovered major immune cell types, but their ability to resolve functionally distinct CD8⁺ TEMRA states varied substantially. Using curated bulk RNA-seq gene sets and Seurat’s ModuleScore() function, we annotated four TEMRA-related cytotoxic subsets: MCTL, DCTL, TCTL, and MAIT. 10X-derived cells showed consistently higher module scores across these subtypes, while Parse-derived cells exhibited negative average enrichment, suggesting reduced capture of signature-defining transcripts (Supplementary Fig 9A and B). This trend translated to subtype distribution as well: over 50% of TEMRA cells in 10X were classified as TCTLs, compared to only 19% in Parse (Supplementary Fig 9C). Our results demonstrate that 10X better preserves the transcriptional identities of CD8⁺ TEMRA subsets, likely due to improved mRNA recovery for short, cytotoxic-associated transcripts.

Mechanistically, these platform differences can be attributed in part to protocol-specific biases. 10X captures 3’ ends of transcripts in unfixed cells, favoring detection of shorter mRNAs and ribosomal genes^23^. Conversely, Parse uses a whole-cell, fixed-cell approach that can diminish recovery of short transcripts. This was especially evident in T cells, where Parse showed reduced expression of canonical markers (*CD3D*, *CD8A*, *CD3G*) and a skew toward NK-like annotations in tools like Azimuth (Supplementary Fig 8). Although Parse detected a broader range of transcript lengths, including longer extracellular matrix and signaling genes, transcriptome counts of cytotoxic T cells were lower compared to 10X (i.e. *GNLY, GZMB*).

Interestingly, both platforms identified core T cell biology, but with different sensitivities. GO enrichment analysis of platform-enriched transcripts revealed stronger enrichment for T cell receptor signaling, cytotoxic effector pathways, and T cell differentiation in 10X data (Fig 5C). These findings are particularly relevant for studying diseases like tuberculosis, where CD8⁺ TEMRA cells play a protective role in latent infection^24^. In contrast, Parse identified more genes associated with structural and matrix-related functions, making it potentially advantageous for profiling tissue-resident or stromal populations.

We also observed that transcript length is a major factor driving detection bias. Genes overexpressed in Parse were significantly longer on average (Fig 5B), supporting the hypothesis that Parse preferentially captures longer transcripts. However, this length bias alone could not explain the misclassification of TEMRA cells as NK cells, suggesting that reduced capture of defining T cell transcripts in Parse is the primary cause of annotation error. Additionally, Azimuth PBMCs references were generated using 10X-derived cells, suggesting the need for more curated cell type references as new transcriptome methods are developed.

While our analysis focused on T cells, particularly CD8⁺ TEMRA populations, we noted discrepancies in other cell types as well, including monocytes. These differences warrant further investigation, especially in the context of lineage-specific gene expression patterns. Additionally, emerging chemistries such as 10X GEM-X v4 and Parse Evercode WT v3 may reduce some of the technical limitations we observed, and future benchmarking studies should include these updated kits.

In conclusion, our side-by-side comparison of 10X and Parse platforms revealed both shared and divergent capabilities in profiling human immune cells. While both platforms quantify global gene expression accurately, 10X demonstrates greater sensitivity in detecting functionally important short transcripts, particularly in CD8⁺ T cell subsets. Researchers should carefully consider platform-specific biases such as ribosomal gene detection, transcript length, and fixation effects when interpreting scRNA-seq results. These findings highlight the value of benchmarking for experimental design and underscore the need for refined reference annotations to improve cell type classification in single-cell genomics.

## MATERIALS AND METHODS

### Ethics statement

This study was conducted according to the principles expressed in the Declaration of Helsinki. The study was performed in accordance with protocols approved by the institutional review board at University of California, Los Angeles. All donors provided written informed consent for the collection of peripheral blood and subsequent analysis (UCLA Institutional Review Board #11-001274).

### Isolation of human Peripheral Blood Mononuclear Cells for scRNA-seq

Peripheral blood mononuclear cells were isolated from the peripheral blood of three healthy donors using a Ficoll-Paque Plus (Cytiva) density sedimentation gradient under endotoxin-free conditions. Isolated PBMCs were subjected to red blood cell lysis using ACK lysis buffer (Thermo Fisher) and then handled as described below.

For 10X Genomics scRNA-seq workflow, cell suspensions were filtered through a 40 µm flowmi cell strainer (SP BEL-ART) and the cell count, and viability were determined using 0.4% trypan blue dye (Gibco). Live PBMCs (viability ≥ 95%) were then resuspended with 1 mL 1X PBS containing 1% UltraPure bovine serum albumin (Thermo Fisher Scientific) and loaded onto the Chip G and Chromium Controller instrument (10x Genomics) to generate gel bead-in-emulsions (GEM) and indexed sequencing libraries according to the Chromium Next GEM Single Cell 3’ Reagent Kits v3.1 user guide. Approximately 30,000 cells per sample were used to achieve the desired cell recovery target of 18,000 cells per sample.

For Parse Biosciences scRNA-seq workflow, cell solutions were passed through a 40 µm Falcon cell strainer (Thermo Fisher) and counted using trypan blue. Live cells (viability ≥ 95%) were then resuspended in cold cell prefixation buffer containing 0.5% BSA Fraction V (Gibco) and fixed using the Evercode cell fixation kit v2 (Parse Biosciences) and stored at -80°C overnight until the start of barcoding and library prep with the Evercode Whole Transcriptome kit v2 according to the manufacturer’s protocol. Approximately 71,000 cells per sample were used to aim for the capture of 18,000 projected barcoded cells per sample.

### Cell sorting of human CD8^+^ memory T cells for scRNA-seq

Terminally differentiated effector memory CD8^+^ T cells re-expressing CD45RA were purified using fluorescence-activated cell sorting (FACS) as previously described^15^. Briefly, cells were labeled with combinations of CD3, CD8, CD45RA, and CCR7 antibodies, and sorted in a BD FACSAria II cell sorter at UCLA flow cytometry core facility to obtain highly purified populations of CD3^+^CD8^+^CD45RA^+^CCR7^-^ TEMRA cells. Zombie Green viability dye (BioLegend) was added to the samples before sorting to exclude nonviable cells. Antibodies used for sorting were as follows: CD3-PerCp (clone SK7, BD Biosciences), CD8-BV605 (clone RPA-T8, BioLegend), CD45RA-APC (clone Hl100, BioLegend) and CCR7-APC-Cy7 (clone G043H7, BioLegend). FACS sorting yielded TEMRA cell populations with ≥ 99% purity.

For 10x Genomics scRNA-seq workflow, cell solutions were handled exactly as described above and barcoded with the Chromium Next GEM Single Cell 3′ Gel Bead Kit v3.1 according to the manufacturer’s protocol.

For Parse Biosciences scRNA-seq workflow, cell suspensions were prepared and fixed as described for PBMCs and then barcoded with the Evercode WT Mini kit v2 (Parse Biosciences) according to the manufacturer’s protocol. Approximately 22,000 cells per sample were used to aim for the capture of 4,000 projected barcoded cells per sample.

### scRNA-seq sequencing, alignment, and data processing

The indexed sequencing libraries from 10x Genomics and Parse Biosciences platforms were subsequently sequenced on an Illumina NovaSeq 6000 S4 flow cell (200 cycles with paired-end reads) at UCLA Broad Stem Cell Research Center (BSCRC) high-throughput sequencing core with average sequencing depth of 20,000 read pairs per cell. Before sequencing the samples, library quality and concentration were measured using the 4200 TapeStation instrument with High Sensitivity D5000 ScreenTape Assay (Agilent) and Qubit Fluorometer with 1X dsDNA HS assay kit (Thermo Fisher) according to the manufacturer’s recommendations.

The resulting FASTQ raw reads were aligned to the GRCh38 human reference genome, and a barcode file, a gene table, and a gene expression matrix were obtained. 10x Genomics scRNA-seq sequencing results were converted from BCL files to FASTQ files using Cell Ranger Single Cell software suite v7. We followed the 10x Genomics Cell Ranger pipeline using default parameters (https://support.10xgenomics.com/single-cell-gene-expression/software/pipelines/latest/using/aggregate). Parse Biosciences scRNA-seq sequencing data were aligned with the Parse Biosciences analysis pipeline v1.1.2 using default parameters.

### scRNA-seq quality control filtering

Using R (v.4), with Seurat package v5^5^, all raw gene expression matrices were downsampled to have the same mean reads per cell using scRecover^22^ (Supplementary Table 3). Doublet cells were calculated using DoubletFinder^23^. We adjusted the doublet prediction rate, for 10X and Parse to be 9.5% and 3%, respectively; all other parameters were used as default. Low-quality cells containing more than 15% mitochondrial genes per cell, less than 500 unique genes per cell, more than 20,000 and less than 1500 UMIs were removed. The data was then normalized using the NormalizeData function with default parameters, the FindVariableFeatures function to select 2,000 genes with the highest variance, and the ScaleData function. Unsupervised clustering for UMAP visualization was performed using RunUMAP, FindNeighbors and FindClusters. ModuleScore was implemented using bulk RNA-seq gene sets^15,22^. The 10X Genomics Single Cell Next GEM 3’ v3.1 and Parse Evercode WT v2 datasets were integrated with the Seurat v5 IntegrateCells (integration=”harmony”)^24^.

### Cell type annotations and gene ontology analyses

Clusters were annotated using RunAzimuth for PBMCs using the ‘pbmcref’ reference and default setting. CD8^+^ TEMRA cells were annotated as mentioned above and with SingleR^25^using the MonacoImmuneData^26^ with default settings and RunAzimuth using the ‘pbmcref’ reference. Gene set enrichment analysis was performed with ClusterProfiler^27,28^ using the significant genes (adjusted p-value < 0.05; avg-log2-FoldChange > 0.5) derived from FindMarkers function using the integrated platform dataset including 10X and Parse cells. The same gene list was used to calculate the gene length using gencode.v32.primary_assembly.annotation.gtf to annotate all protein coding genes.

### Bulk RNA-seq analyzes

Raw bulk RNA-seq counts were collected from GSE159337^19^ using only healthy samples and filtered with a minimum RNA expression of 5. Gene lengths were annotated as previously described.

## Data Availability

All single-cell RNA-seq is available to download from Gene Expression Ominibus (GSE285843).

## Author Contributions

Conceptualization: AE, BJAS; Funding Acquisition: MP; Investigation: AE, BJAS; Methodology: AE, BJAS; Supervision: MP; Visualization: AE, BJAS; Writing – Original Draft Preparation: AE, BJAS; Writing – Review Editing: MP, AE, BJAS.

## Acknowledgements

We thank I. Williams and the UCLA JCCC Flow Cytometry Core Facility for assistance with FACS sorting; and S. Feng and the UCLA Broad Stem Cell Research Center (BSCRC) high-throughput sequencing core for assistance with Illumina sequencing. We thank the Robert L. Modlin and their lab for equipment and guidance.

